# Higher order synthetic lethals are keys to minimize cancer treatment effects on non-tumor cells

**DOI:** 10.1101/2025.01.31.635848

**Authors:** Mehdi Dehghan Manshadi, Payam Setoodeh, Amin Ramezani, Amin Reza Rajabzadeh, Habil Zare

## Abstract

Metabolic rewiring in cancer cells facilitates the provision of essential precursors for the unbridled growth of tumors. Exploring these cancer-specific metabolic changes offers potential selective therapeutic strategies. However, targeting a single essential gene in cancer treatment often faces challenges, including resistance, lack of targetable oncogenes, and potential harm to non-tumor cells. Attacking multiple targets is hypothesized as a solution to overcome these issues, e.g., a synthetic lethal set, defined as a minimal combination of non-lethal genetic mutations leading to cell death. This study examined the potential of synthetic lethal sets to identify selective drug targets for 13 cancers and the corresponding non-tumor tissues, utilizing context-specific genome-scale metabolic models. To ensure the minimization of therapeutic side effects, this work introduced the concept of *strictly-selective* drug targets (SSDTs) and the harmlessness of identified targets in all 13 different non-tumor tissues was meticulously verified. Accordingly, for 13 types of cancers, over 500 SSDTs were identified, predominantly including higher-order synthetic lethal sets with more than two targets in each set. Interestingly, for specific cancers where single essential or synthetic lethal genes could not provide acceptable solutions, SSDTs were provided by higher-order synthetic lethal sets. Therefore, for the first time, this study successfully showed that leveraging higher-order synthetic lethal sets holds the key to promising strictly-selective solutions. Furthermore, nine quadruple SSDTs were identified which commonly target five different cancers without harming any of the 13 non-tumor tissues. Further experimental validation of these findings is required to select the most promising treatments for clinical studies.

## Introduction

Metabolic rewiring leads to distinct changes in the flux rates through various pathways in cancer cells. Otto Warburg was the first to highlight these metabolic variations, revealing that cancer cells prefer aerobic glycolysis over oxidative phosphorylation (1). This phenomenon, known as the Warburg effect, illustrates how these metabolic alterations in cancer cells support the increased demand for fatty acids, nucleotides, and amino acids (2). Extensive research has been conducted on aerobic glycolysis to better understand cancer cell metabolism (3–13).

In addition to the Warburg effect, various other metabolic alterations have been reported in different cancers. One such alteration is the up-regulation of the Glutaminolysis pathway in cancer cells, which collaborates with glycolysis up-regulation to supply both energy and precursors, supporting higher proliferation rates (14–16). Furthermore, the literature indicates the deregulation of other pathways, such as the lipid synthesis pathway (17), the branched- chain amino acid metabolism, the serine synthesis pathway (18), and the pentose phosphate pathway (19). Metabolic rewiring in cancer cells also presents unique opportunities for immune evasion (20), apoptosis disruption (21), and the utilization of conventional waste products from cells, including ammonia, ketone bodies, acetate, and lactate (18).

Characterization and identification of the mentioned cancer-specific metabolic alterations pave the way for developing selective therapeutic strategies that exclusively affect cancer cells without harming non-tumor cells (22–24). Because genome-scale metabolic models (GEMs) are known as accurate mathematical representations of metabolism and promising tools for studying metabolic rewiring strategies at a cell-wide level (25–27), different computational approaches have been developed to use these models for identifying metabolic cancer drug targets (28, 29). Folger et al. (30) provided one of the first studies in the field of drug target identification using GEMs. The authors provided a single model using expression data from different cancers and found 52 drug targets, of which 21 were targeted by experimental, approved, or known anticancer drugs. To generate context- or patient-specific GEMs, different approaches have been developed and applied to identify different cancer therapeutics (31). Yizhak et al. (32) built more than 280 healthy and cancerous models using PRIME (personalized reconstruction of metabolic models). They utilized this set of cell-specific models to predict selective drug targets, experimentally validating the top predicted target, *MLYCD*, and investigating the depletion effects of this gene. In another study, Gatto et al. (33) used the INIT algorithm (34) to generate GEMs for clear cell renal cell carcinoma (ccRCC) and successfully identified five metabolic genes as selective drug targets, predicted to be dispensable in non-tumor cell metabolism. Barrena et al. (35) extended the gMCS approach by integrating linear regulatory pathways with metabolic models using Human1 and regulatory databases. They identified new essential genes and synthetic lethal pairs in cancer cell lines. Larsson et al. (36) used a generic GEM to identify five selective essential genes for glioblastoma, with *in vitro* or *in vivo* evidence of the essentiality of 4 out of the 5 were reported in the literature. Pacheco et al. (37) introduced the rFASTCORMICS algorithms and built 10,005 GEMs for 13 different types of cancers. The authors performed a single gene essentiality analysis to identify selective drug targets and used colorectal cancer as a successful experimental test case for drug repurposing of mimosine, ketoconazole and naftifin.

Despite valuable single-drug targets being identified using the mentioned approaches, based on the literature, targeting only one gene can lead to various difficulties (38). The survival dependency of a tumor on an oncogene or oncogenic pathway is a phenomenon known as oncogene addiction. Currently, this phenomenon forms the basis of most genotype-targeted cancer therapeutics. However, gain-of-function oncogenes are not targetable in all tumors, resulting in common resistance to the administered therapy (39). Furthermore, identifying unmarked oncogenes, the cancer-specific driver genes with no evidence of genetic alterations is challenging. To overcome these challenges, scientists suggest using the synthetic lethality principle (39–41). Synthetic lethality describes a gene pair with viable mutations of each gene separately, while their simultaneous mutation is deadly. Additionally, it is hypothesized that it is possible to identify selective therapeutic strategies by targeting a synthetic lethal (SL) gene belonging to a cancer-relevant mutation (42). Furthermore, it is expected that SL sets offer new therapeutic approaches by targeting cancers with pathway alterations that are generally regarded as undruggable (43). Although the capabilities of SLs for providing selective drug targets are mentioned in the literature, limited studies were conducted for the identification of these drug targets. In this context, Frezza et al. (44) conducted a study on fumarate hydratase Fh1-deficient kidney cells. The researchers created context-specific genome-scale metabolic models (GEMs) for cell lines with and without Fh1, using gene expression data. They predicted gene knockouts were lethal to Fh1-deficient cells but did not affect non-tumor cells. The analysis highlighted reactions, primarily in the heme metabolism pathway, as synthetically lethal with Fh1 deletion. In another study, Zhang et al. (45) performed an exhaustive search to identify 44 double and 95 triple-selective drug targets for hepatocellular carcinoma (HCC) using a generic GEM from Argen et al. (46).

Although synthetic lethality is assumed to be a more reliable approach to combat cancers through selective drug targets compared to attacking single essential genes, to the best of the authors’ knowledge, no systematic study has been conducted to demonstrate this advantage. Therefore, this work aims to identify SL gene sets predicted by context-specific genome-scale metabolic models to determine the capabilities of the concept of synthetic lethality for the identification of *strictly-selective* drug targets. Here, the term “*strictly-selective*” refers to drug targets that affect cancer cells while causing minimal damage not only to the non-tumor tissue of the corresponding cancer but also to non-tumor tissues of other organs. To achieve this goal, context-specific models were created for 13 different pairs of cancerous and non-tumor tissues using the rFASTCORMICS algorithm. The names and abbreviations of the studied cancers are listed in Table 1.

**Table 1.** The TCGA abbreviations of the cancers studied in this paper.

| TCGA name | Expanded name |
| --- | --- |
| COAD | Colon adenocarcinoma |
| BRCA | Bladder Urothelial carcinoma |
| LUAD | Lung adenocarcinoma |
| LIHC | Liver hepatocellular carcinoma |
| LUSC | Lung squamous cell carcinoma |
| UCEC | Uterine Corpus endometrial carcinoma |
| HNSC | Head and neck squamous cell carcinoma |
| STAD | Stomach adenocarcinoma |
| KIRC | Kidney renal clear cell carcinoma |
| PRAD | Prostate adenocarcinoma |
| KICH | Kidney chromophobe |
| THCA | Thyroid carcinoma |
| KIRP | Kidney renal papillary cell carcinoma |

For these models, essential genes, SLs, and higher-order SLs, each comprising up to four genes, were identified. Subsequently, potential *strictly-selective* drug targets were determined. The results showed that when a few or even no essential genes can be targeted due to causing side effects for different non-tumor cells in the human body, higher-order SL sets are the keys to *strictly-selective* drug targets that discriminate solely against cancerous tissue. Accordingly, in addition to the identification of several *strictly-selective* potential drug targets for 13 cancers, for the first time, this study successfully showed the capabilities of the concept of synthetic lethality for providing strictly selective drug targets for cancers using genome-scale metabolic modeling and *in-silico* experimentation.

## Methods

### Reconstruction of context-specific model

In this study, 26 context-specific models, consisting of 13 generic control models and 13 generic cancer models, were reconstructed using rFASTCORMICS (37), as a promising tool to generate accurate models capable of predicting the outcome of gene knockout strategies. The Recon 2.04 (47) genome-scale reconstruction (RRID:SCR_006345) was chosen as the reference model and the RNA-seq data (GSE62944 (48)) were collected from the TCGA Research Network (http://cancergenome.nih.gov/). The assignment of samples to cancerous and non-tumor tissues was based on their pathology information available from TCGA. This process could not be randomized. Also, power analysis was not relevant here because we used all existing data.

While arbitrary thresholds for RNA-seq data can significantly impact model precision in tracking expressed and non-expressed genes (49), rFASTCORMICS adopts an approach proposed by Hart et al. (50) to determine active genes. Specifically, rFASTCORMICS generates a density plot for each sample based on its log2-transformed FPKM (Fragments per Kilobase of transcript per Million mapped reads) values. Subsequently, a Gaussian curve is fitted to the right half of the main peak (expression curve). The mean value of this curve is utilized as the expression threshold. Then, the obtained curve is subtracted from the density curve, and another Gaussian curve is fitted to this part of the density curve (inexpression curve).

The rFASTCORMICS uses the mean value (μ) and standard deviation (σ) of the right-hand Gaussian curve to convert log2-transformed FPKM values to the zFPKM values.

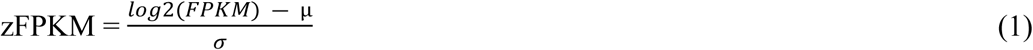

The mean of the inexpression curve was chosen as a threshold if it was above -3 zFPKM; otherwise, -3 was taken, and genes with lower zFPKM values were considered inactive with a score of -1. Conversely, 0 zFPKM was chosen as the expression threshold, and genes with higher zFPKM values were considered active with a score of +1. Genes with zFPKM values between these two thresholds were considered to have an *unknown expression status* with a score of 0. Finally, using gene-protein-reaction (GPR) rules, expression scores for genes were mapped to corresponding reactions, and bounds for inactive reactions were set to zero.

### Extracting SL sets

Extracting SL gene pairs from reconstructed models is not computationally challenging. However, extending the synthetic lethality principle to gene sets with more than two genes and attempting to identify higher-order SL sets (HOSLs) presents significant computational challenges. As the number of genes in the SL set increases, the corresponding search space grows exponentially, making the search process extremely time-consuming. Therefore, an algorithm is required to explore this expanding search space efficiently. In this study, Rapid-SL (51) was applied to search for HOSLs up to quadruples (four genes in a set) for 13 cancerous models. Alternative methods for determining all HOSLs, including gMCS (52) and Fast-SL (53) which might not be practically as fast as Rapid-SL.

### Strictly-selectivity criteria

The primary goal of this work was to identify *strictly-selective* potential drug targets. In this study, *strictly-selective* drug targets refer to essential genes or SL gene sets. These targets aim to reduce the growth rate of cancer cells while leaving all non-tumor tissues of other organs in the human body unaffected in terms of growth rate reduction. To mathematically describe these criteria, certain thresholds were used. Here, a solution was considered lethal if its knockout reduced the growth rate of cancer cells by over 50% (37). Furthermore, not affecting non-tumor tissues was defined as causing less than a 10% (37) reduction in the growth rate of non-tumor tissues when the gene or gene set was knocked out from the corresponding models. Figure 1 shows an overview of the procedure considered in this work.

**Figure 1.**
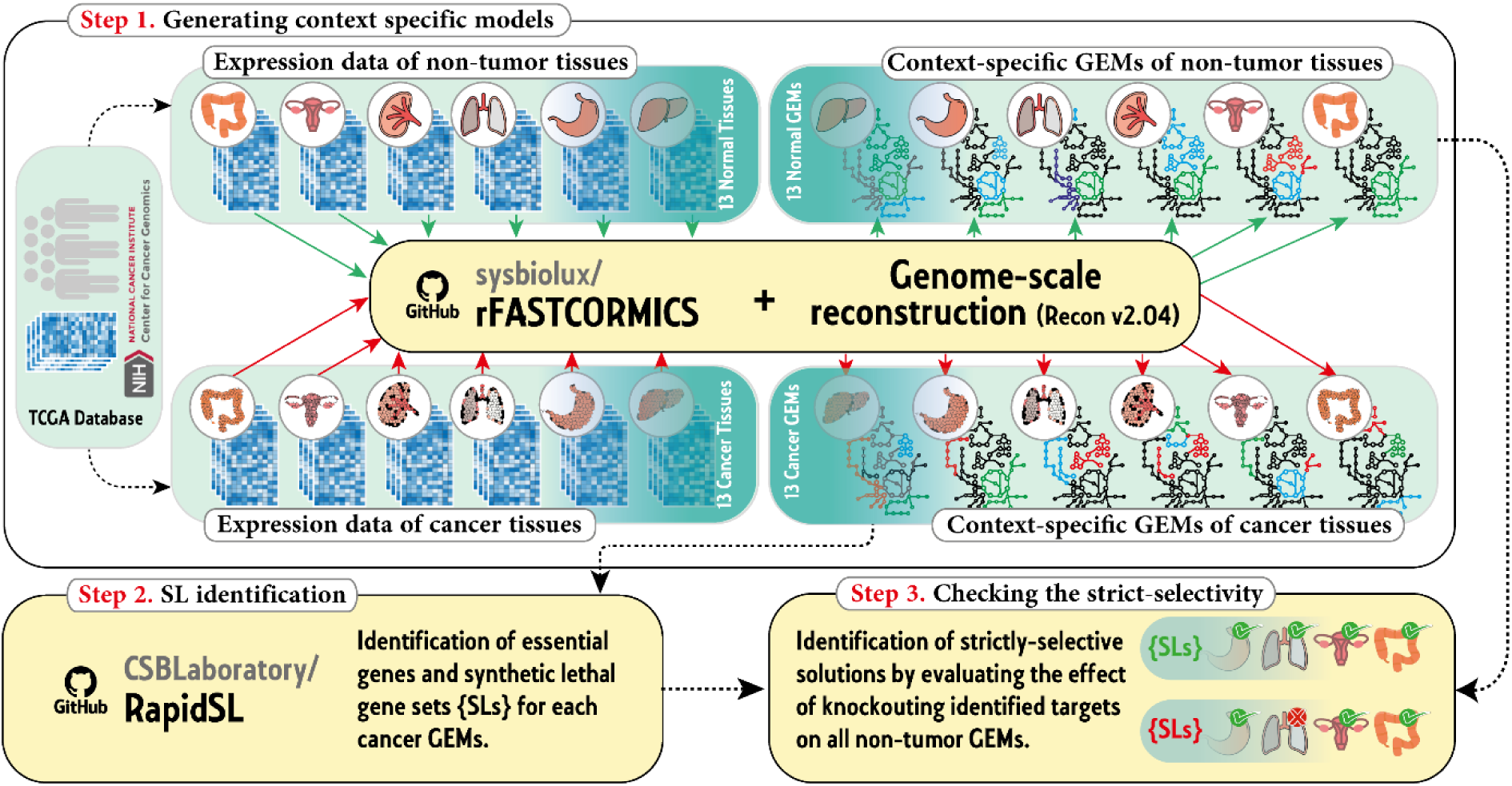
An overview of the procedure considered in this study.

## Results

This study identified essential genes and SL gene sets for 13 context-specific genome-scale models of different cancers. Additionally, besides identifying SL genes, this study successfully obtained triple and quadruple synthetic lethal gene sets, referred to as higher-order SL sets. The list of *strictly-selective* drug targets obtained in this work is reported in supplementary file S1.

### Cases with essential genes as *strictly-selective* solutions

Here, the reported results are explained using colon cancer as an example. The number of essential genes and synthetic lethal sets obtained for the COAD GEM is reported in Table 1.

This table illustrates how the number of solutions decreased when their selectivity was tested using the GEM of the non-tumor tissue of the colon.

**Table 1.** The number of essential and SL solutions obtained for colon cancer.

|  | Single<br>essentials | Double<br>SLs | Triple<br>SLs | Quadruple<br>SLs |
| --- | --- | --- | --- | --- |
| **Lethal for cancer cells** | 76 | 64 | 145 | 416 |
| **Selective for non-tumor colon cells** | 12 | 26 | 57 | 135 |
| **Percentage of selective solutions (%)** | 15.8 | 40.1 | 39.3 | 32.5 |

Although the solutions obtained in this step are potentially selective for cancerous colon tissue cells, there is still a chance of damaging other non-tumor cells. Therefore, the effect of the remaining solutions on other non-tumor tissues was examined step by step by considering the selectivity criteria for other non-tumor cells such as breast, lung, etc. By considering each additional non-tumor tissue, some potential solutions may be found to be non-selective and omitted from the list of potential selective drug targets. When considering all 13 non-tumor tissues, many solutions faded away. As a result, *SLC38A3*, a member of the transport of inorganic cations/anions and amino acids/oligopeptides pathway, has been identified as a single *strictly-*selective drug target. In comparison to *SQLE* proposed by Pacheco et al. (37), which has side effects on other non-tumor tissues such as the liver or kidney, *SLC38A3* is expected to have minimal side effects on all 13 non-tumor tissues. Asparagine (DB00174) is one of the drugs that inhibits *SLC38A3* and can be considered a candidate for validating this *strictly- selective* drug target.

Although no double *strictly-selective* SL was found, two three-membered synthetic lethal sets were identified for COAD. The first set includes *RPE*, *HSD11B2*, and *G6PD*. Two of these genes also appear in the second synthetic lethal set, which includes *RPE*, *HSD11B2*, and *PGLS*.

These genes are functionally related. Specifically, *RPE*, *G6PD*, and *PGLS* contribute to the Pentose phosphate pathway and the gene *HSD11B2* is related to the Glucocorticoid biosynthesis pathway. Various drugs are reported to have effects on *G6PD*, *PGLS*, and *HSD11B2* (see Supplementary file S4); however, *RPE* is not listed as a target for any drugs on DrugBank (54). Because all genes in the set of *RPE*, *G6PD*, and *HSD11B2* are reported to have fold changes greater than 1 (55), we propose this lethal set as a suitable candidate for further investigations.

By increasing the cardinality of the SLs to four, the number of strictly-selective drug targets increases to 9 sets. The genes of these nine sets are depicted in Figure 2. The characteristics of these genes are investigated in the discussion section.

**Figure 2.**
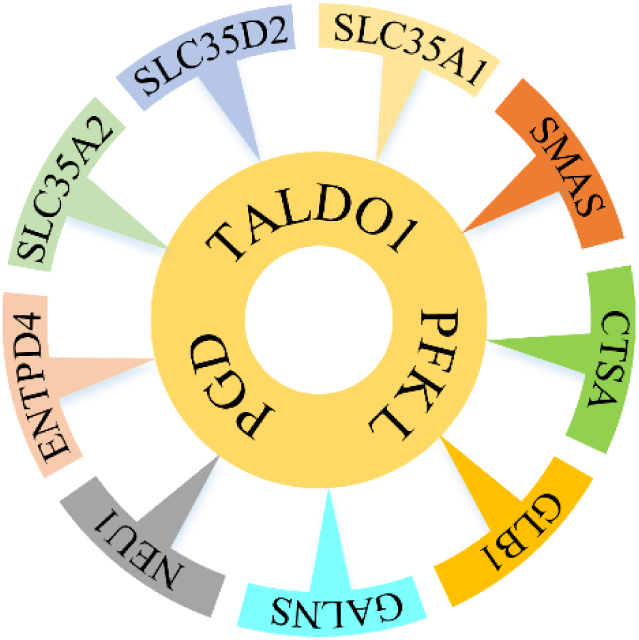
The nine mutual quadruple synthetic lethal sets among COAD, BRCA, HNSC, UCEC, and STAD. All sets have three genes *PFKL*, *TALDO1*, and *PGD,* in common.

To find out how protecting further non-tumor tissues would change the number of potential solutions, the number of single essential, double SL, triple SL, and quadruple SL solutions that remained selective at each step is presented in Figure 3.

**Figure 3.**
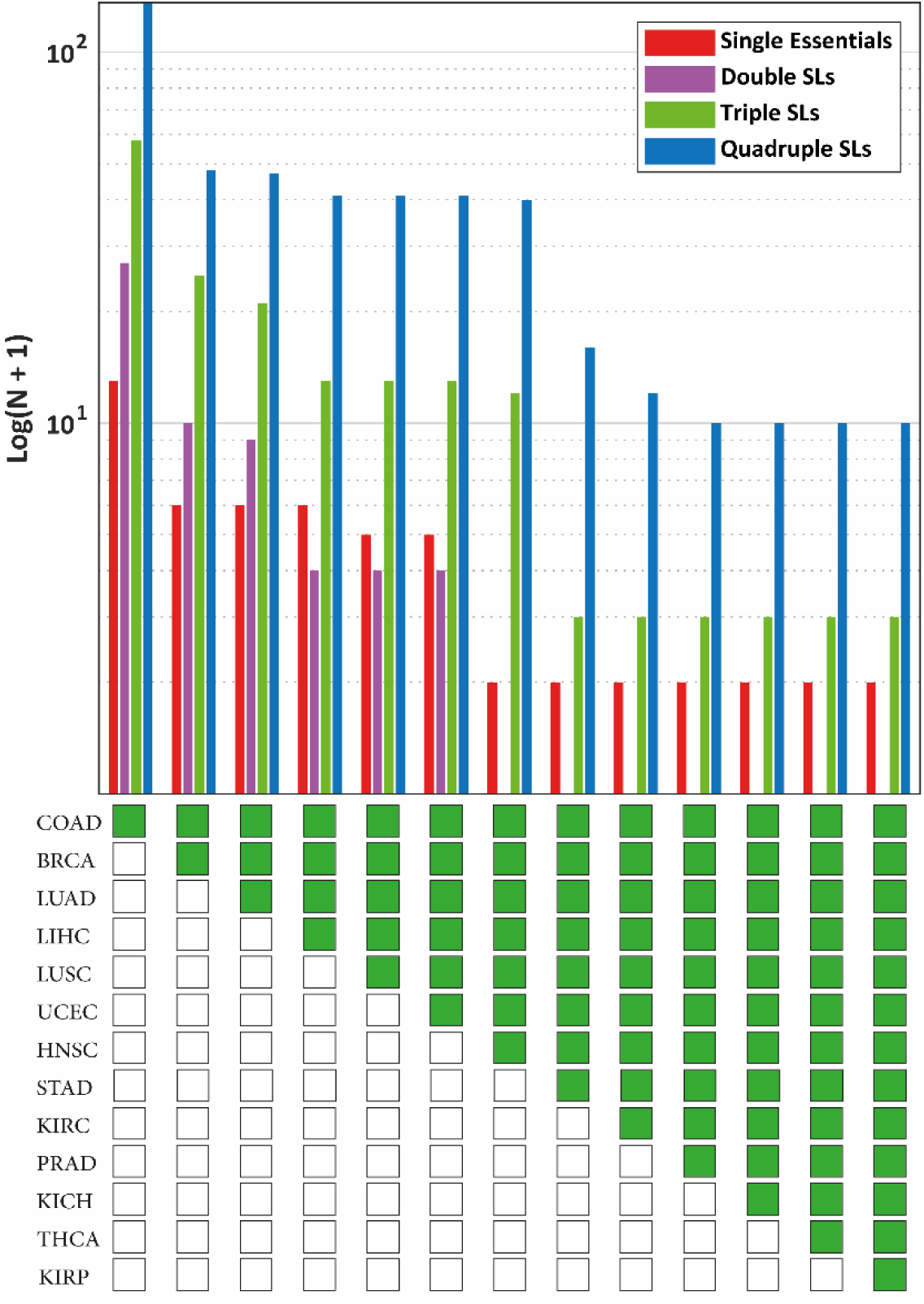
The number of potential selective drug targets identified by considering different non-tumor cells as protected and COAD as the target. Considered non-tumor tissues for each column are marked in green below the chart.

In Figure 3, from left to right, as the number of tissues considered protected increases, the number of selective drug targets decreases. Considering only six non-tumor tissues (including the non-tumor tissue of the colon itself), all double SL sets were disqualified, leaving only one essential gene as a selective target. However, 11 higher-order synthetic lethal sets (nine quadruple SLs and two triple SLs) remained as *strictly-selective* solutions.

As an example of these higher-order SLs, the simultaneous knockout of *PFKL*, *TALDO1*, *PGD*, and *CMAS* was identified as deleterious for COAD cells, with potentially minimal side effects on the 13 non-tumor tissues. For other cancers, the same figures were generated and reported in Figure 4 and Figure 5.

**Figure 4.**
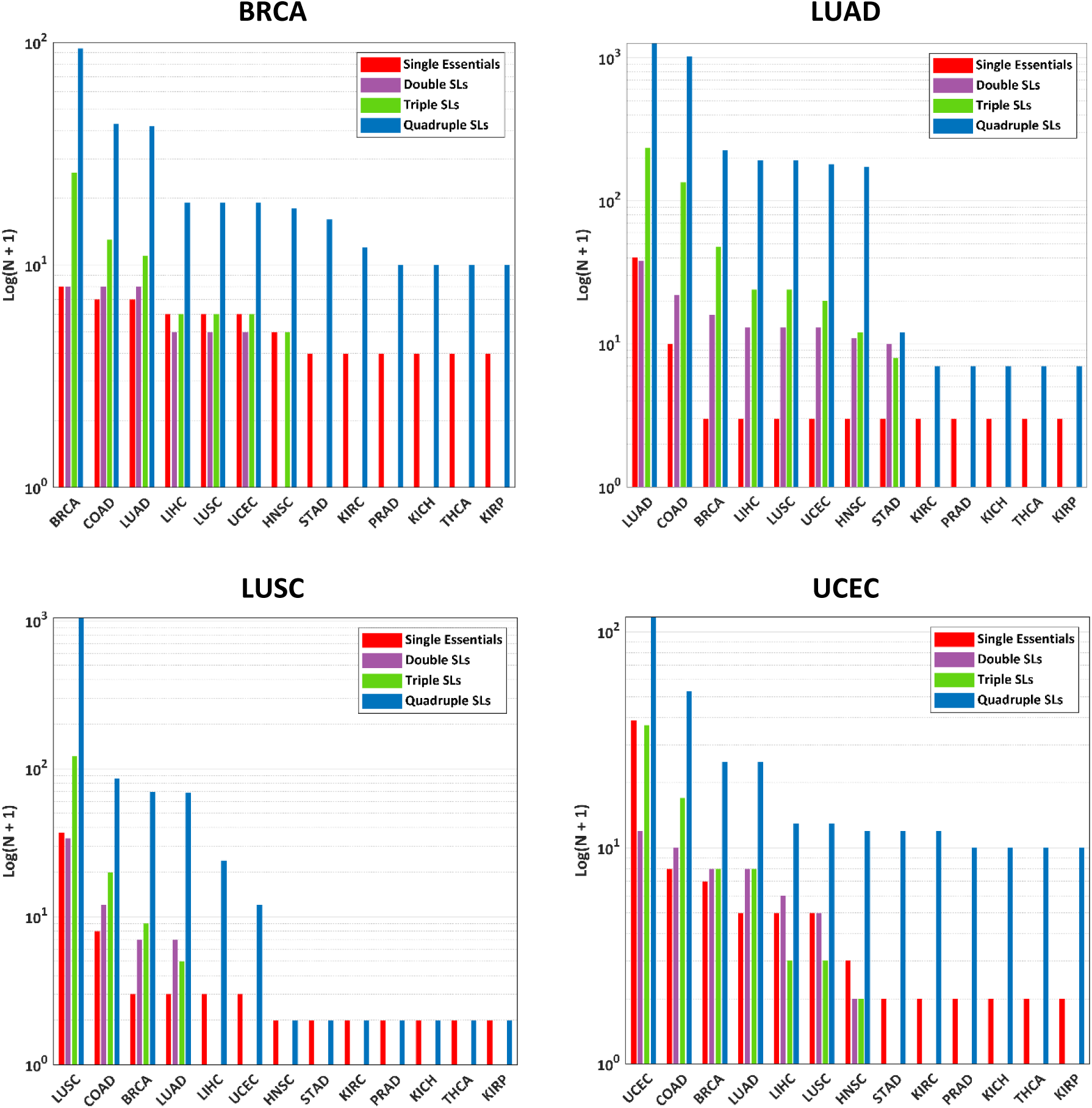
The number of potential selective drug targets identified by considering different non-tumor cells as protected and BRCA, LUAD, LUSC, and UCEC as the targets.

**Figure 5.**
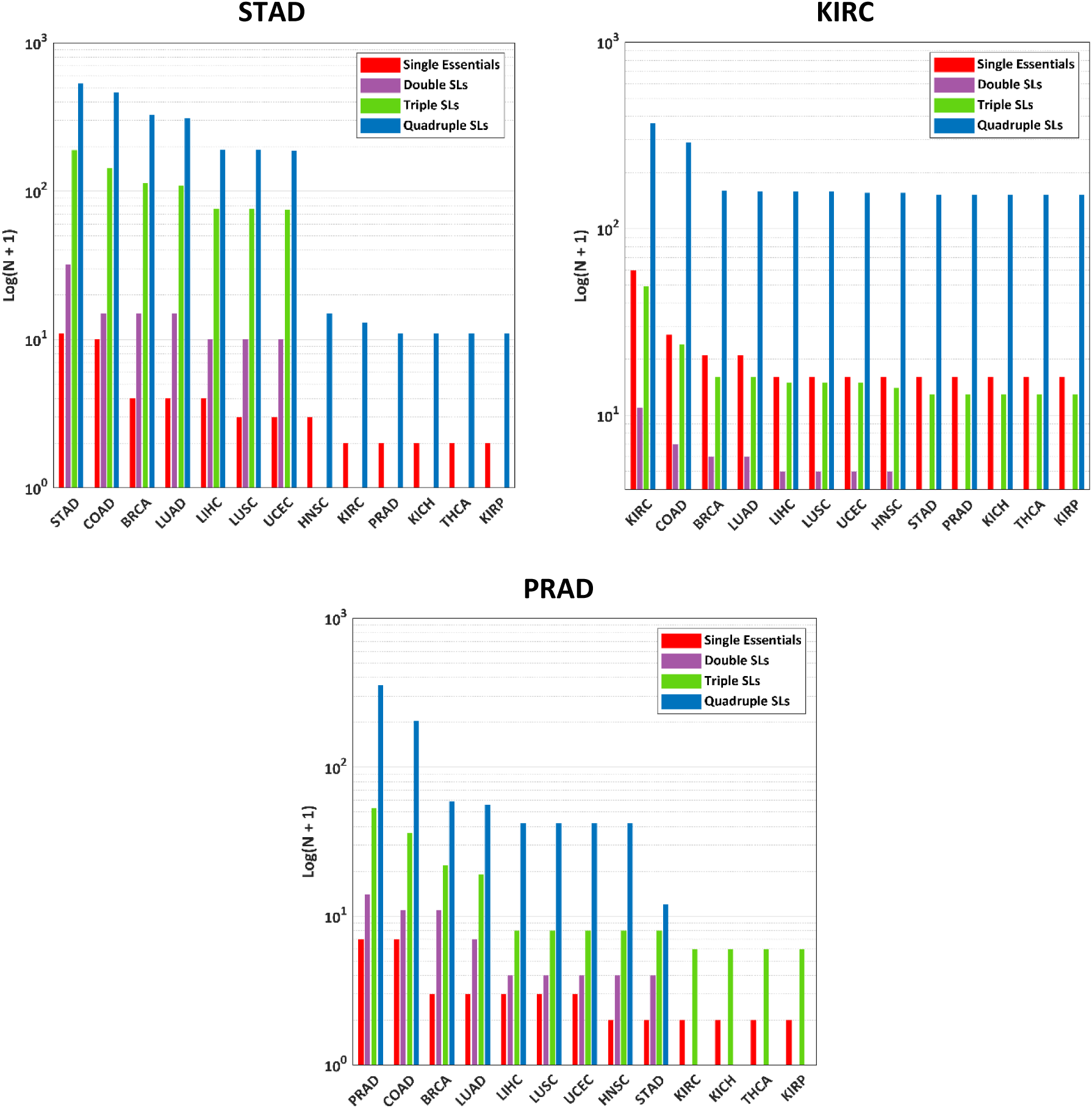
The number of potential selective drug targets identified by considering different non-tumor cells as protected and STAD, KIRC, and PRAD as the targets.

For these eight cases (shown in Figure 3 to Figure 5), no synthetic lethal set (sets with two targets) was found to meet the *strictly-selectivity* criteria, while higher-order synthetic lethal sets, especially quadruple SLs, showed their dominance in offering potential solutions.

### Cases without any essential genes as *strictly-selective* solutions

After obtaining *strictly-selective* drug targets for all 13 cancers, it was revealed that there is no single essential drug target for five cases LIHC, HNSC, KICH, THCA, and KIRP. This suggests that essential genes of these cancers are also essential for at least one non-tumor tissue. Interestingly, for all of these cases, SLs, and more significantly, higher-order synthetic lethals, provide acceptable solutions. The number of selective solutions at each step for these five cases is shown in Figure 6.

**Figure 6.**
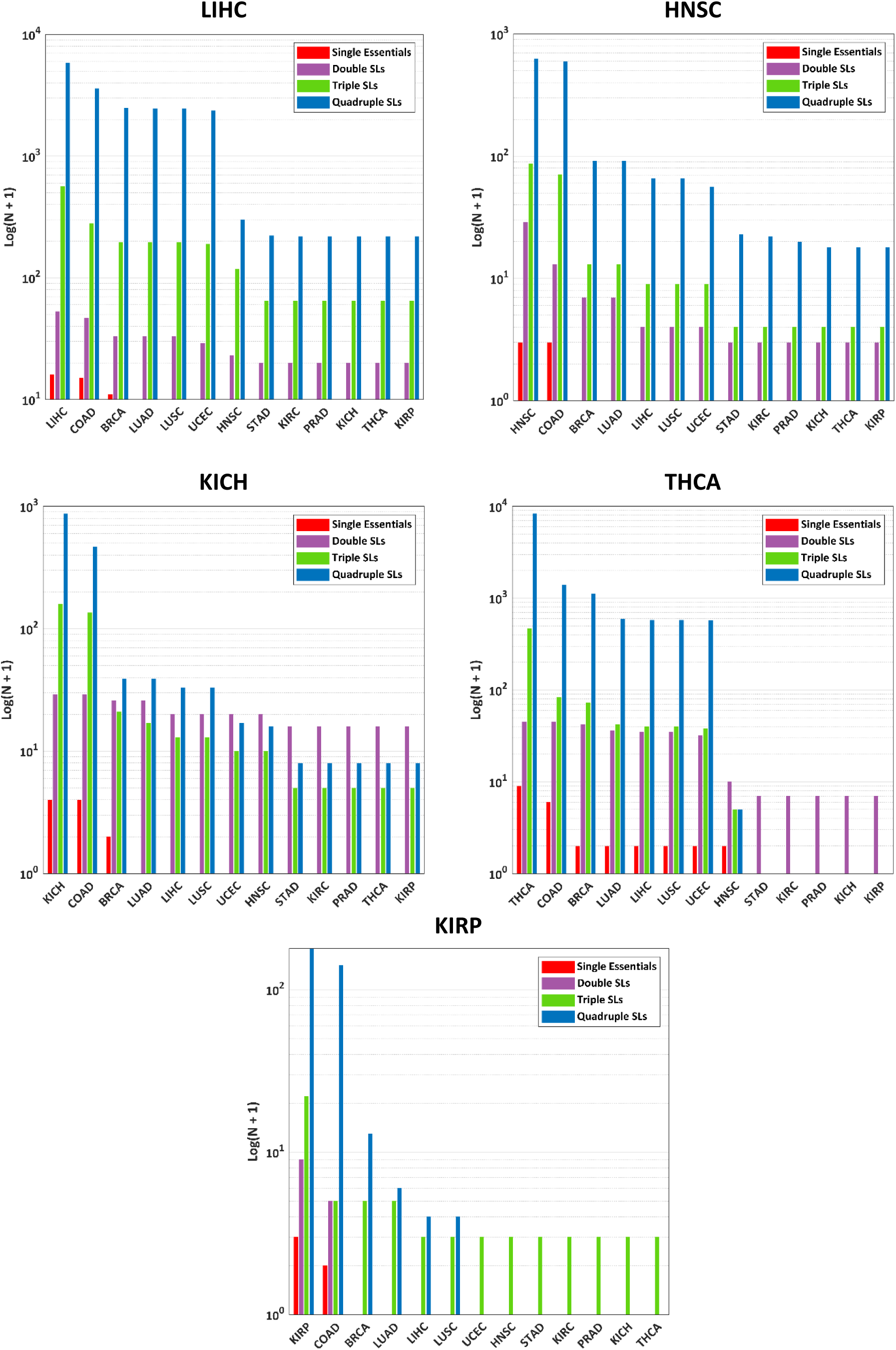
The number of potential selective drug targets identified by considering different non-tumor cells as protected and LIHC, HNSC, KICH, THCA, and KIRP as the targets.

**Figure 7.**
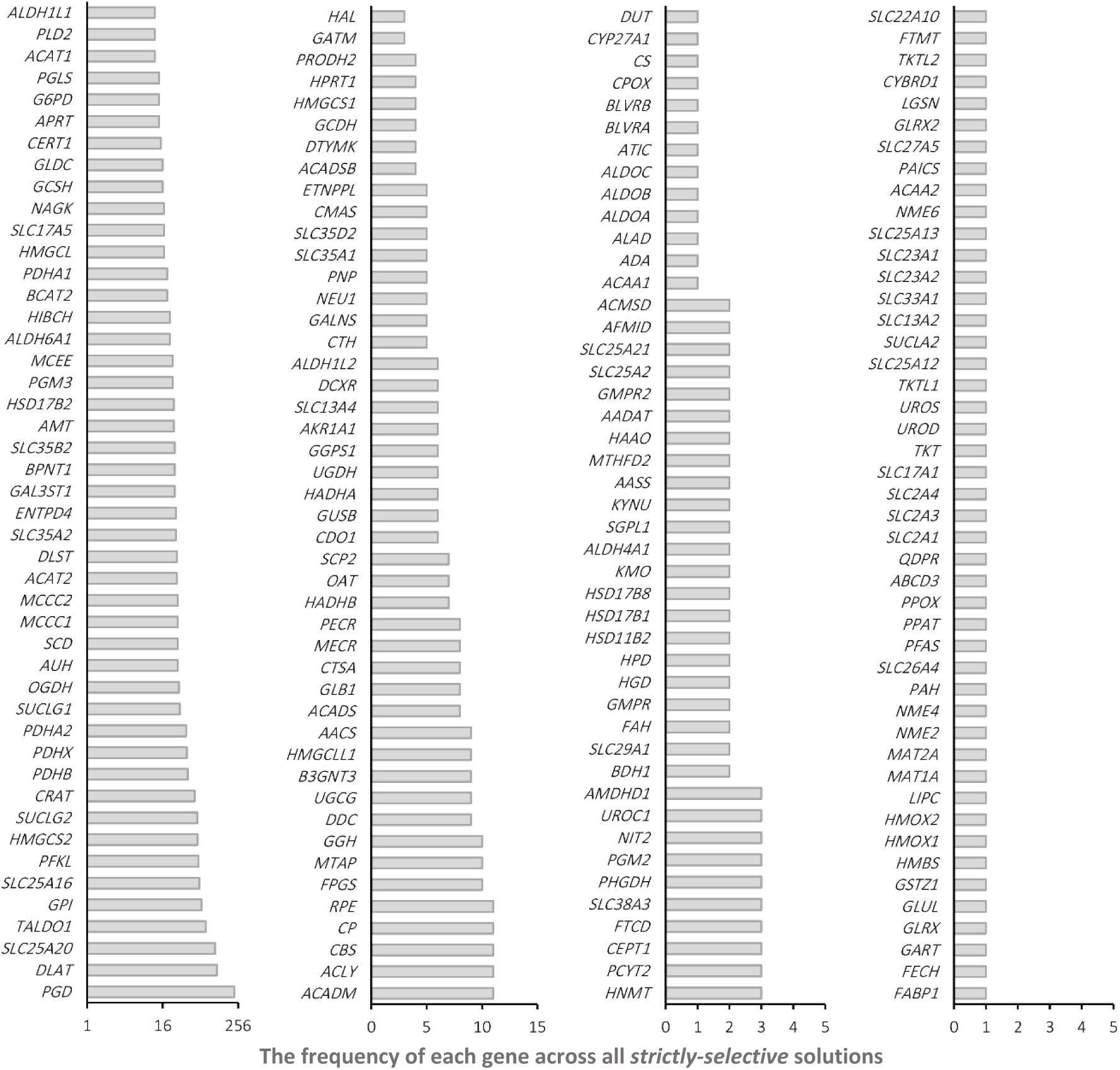
The frequency of different genes across all obtained solutions. This figure shows how many times a gene was identified as a target across all single essential and synthetic lethal solutions.

### Exploring obtained solutions for different cancers

In this section, different aspects of the obtained solutions are investigated. By considering all obtained solutions, the most mentioned gene, as shown in Figure 6, is Phosphogluconate Dehydrogenase (*PGD*), followed by Dihydrolipoyl transacetylase (*DLAT*) and solute carrier family 25 member 20 (*SLC25A20*). Other important genes, such as Transaldolase 1 (*TALDO1*) and 6-phosphofructokinase liver type (*PFKL*), were targeted in several solutions.

The relationship between different genes in obtained solutions is visualized in Figure 8 based on their co-presence in each SL set. Because various solutions were identified for LIHC and KIRC, their graph is not presented here (see supplementary file S2).

**Figure 8.**
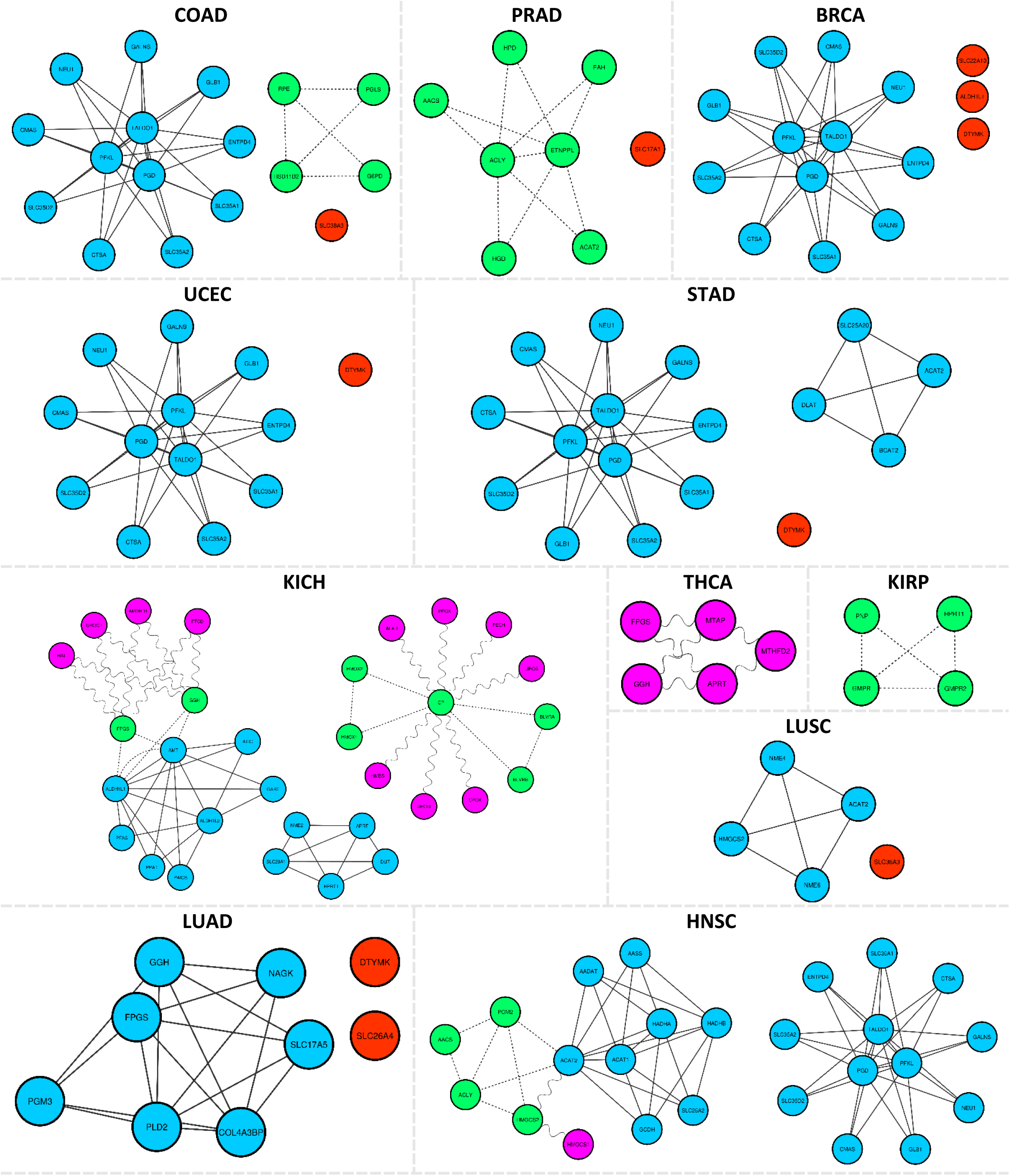
The relation between different genes of identified *strictly-selective* solutions. Single essentials are shown in red, double SLs in purple and connected by wavy lines, triple SLs in green and connected by dashed lines, and quadruple SLs in blue and connected by solid lines.

According to Figure 8, COAD, BRCA, HNSC, UCEC, and STAD share a group of nine quadruple SL sets that consist of a core of genes, including *PFKL*, *TALDO1*, and *PGD.* In each set, the fourth gene acts as a final shot and reduces the growth rate of the *ΔPFKL,ΔTALDO1,ΔPGD* mutant to less than 50% of the corresponding cancer cell.

## Discussion

In this work, *strictly-selective* potential drug targets were obtained by identifying those that are lethal for cancerous cells while inactive, not only in the healthy tissue of the targeted cancer, but also in the other non-tumor tissues of the human body. Although for selective targeting of cancer cells, it is a common practice to necessarily target a cancer-specific mutant gene alongside a non-essential gene (39, 40), this study did not limit the identification of synthetic lethal pairs to cancer-specific mutations. Through this unbiased approach, a wider range of solutions were identified, which does include solutions based on the cancer specific mutations. Note that the provided results are solely based on in silico experiments, and further examinations are crucial for each specific potential case, especially because some of the obtained solutions, such as *SLC38A3*, *RPE*, *HSD11B2*, or *PGLS* identified as a target for different cancers in this study, have been reported as deleterious in mice (56).

To obtain the *strictly-selective* drug targets, 26 context-specific genome-scale metabolic models were reconstructed for non-tumor and cancerous cells of 13 tissues using the rFASTCORMICS algorithm. Afterward, synthetic lethality analysis on the 13 cancerous models was performed using Rapid-SL to identify double, triple, and quadruple Synthetic lethal gene sets, as well as single essential genes. As a result, desired drug targets were identified for each cancer by checking the *strictly-selectivity* criteria on each essential gene or synthetic lethal gene set using the 13 non-tumor GEMs.

The superiority of higher-order synthetic lethal sets to provide potential solutions was demonstrated by analyzing obtained solutions. It also revealed that no *strictly-selective* solution could be provided by single essential genes for LIHC, HNSC, KICH, THCA, and KIRP, and only synthetic and higher-order synthetic lethal sets were able to offer qualified solutions for these cases. Furthermore, nine quadruple synthetic lethal sets were identified as *strictly- selective* for COAD, BRCA, HNSC, UCEC, and STAD. These nine sets include three shared genes *PFKL* (Phosphofructokinase), *TALDO1* (transaldolase), and *PGD* (Phosphogluconate Dehydrogenase), and nine different genes for each set. The three shared genes are among the most frequently identified genes in the solutions and are involved in both glycolysis and the pentose phosphate pathway, both of which are highly implicated in the development of cancers. Due to the crucial role of pentose phosphate pathway overexpression in supporting the anabolic needs and addressing oxidative stress in glycolytic cancer cells (57, 58), simultaneously targeting both pathways is expected to be effective against cancers.

Phosphofructokinase is an enzyme of the Glycolysis and Gluconeogenesis pathway, and its inhibition has been reported to be effective in suppressing colorectal cancer (59). Additionally, it is known to halt the progression of hepatocellular carcinoma (60) and manage lung cancer (61). Transaldolase is an enzyme related to the pentose phosphate pathway and has been associated with the development of cancer (62). This enzyme plays a vital role in the pentose phosphate pathway within fast-dividing cancer cells, serving as a crucial mediator that enables glycolytic cancer cells to meet their anabolic needs and effectively cope with oxidative stress (58). Additionally, this enzyme is associated with bladder (63), liver (64, 65), and breast (66) cancer. The association between higher Transaldolase expression and decreased responsiveness to *HER2* inhibitors in breast cancer patients suggests a potential role of Transaldolase in cancer drug resistance. Therefore, suppressing Transaldolase as a drug target increases the susceptibility of *HER2*-amplified cell lines to *HER2* inhibition (67).

Phosphogluconate dehydrogenase is one of the pentose phosphate pathway enzymes that has recently gained attention due to its crucial role in the development of tumors and maintaining cellular redox balance (68). Phosphogluconate dehydrogenase is frequently overexpressed in various types of cancer, promoting the ability to grow and spread in different cancer cells, such as breast (69, 70), ovarian (71), and lung (72). Additionally, overexpression of Phosphogluconate dehydrogenase is related to the development of resistance to radiotherapy (73) and chemotherapy (68, 74, 75) in different cancers.

Besides the three shared genes, nine distinct genes exist in each of the nine identified *strictly- selective* potential drug targets. These genes are related to five pathways: *CTSA*, *GLB1*, and *NEU1* are related to the sphingolipid metabolism pathway, which is altered in the development and progression of different cancers, especially colon cancer (76–81). *SLC35A1*, *SLC35A2*, and *SLC35D2* are related to the solute carrier family 35, which belongs to the transport pathway of the Golgi apparatus. Furthermore, the SLC family has been reported to have a significant role in anticancer drug resistance (82). *SLC35A2* has been reported to be upregulated in many cancers (83, 84), including colon cancer. *CMAS*, or Cytidine monophosphate N- acetylneuraminic acid synthetase, is related to amino-sugar metabolism. Knockdown of *CMAS* has anticancer activity in triple-negative breast cancer (85). *GALNS*, or N-acetylgalactosamine- 6-sulfatase, is related to keratan sulfate biosynthesis and is reported to have higher expression in many cancers (86). *ENTPD4*, or ectonucleoside triphosphate diphosphohydrolase 4, is known to have hydrolase activity and specifically functions as a UDP phosphatase, being related to nucleotide metabolism. However, *ENTPD4* is a novel gene that has been investigated in a few research studies. Recently, a study showed that *ENTPD4* is overexpressed in gastric cancer tissues (87). Hence, its downregulation or knockout may provide a path for competing cancers.

## Conclusion

In this work, for the first time, the superior capability of higher-order synthetic lethal sets over single essential genes in providing *strictly-selective* drug targets was demonstrated using genome-scale metabolic models. Besides reporting over 500 solutions identified as strictly- selective for 13 types of cancers, nine quadruple SLs are specifically investigated as common solutions between five cancers. Attacking any of these nine SLs results in the disruption of glycolysis and the pentose phosphate pathway along with one of the sphingolipid metabolism, solute carrier family, amino-sugar metabolism, keratan sulfate biosynthesis, or nucleotide metabolism pathways. This simultaneous attack suggests a key to effectively and selectively targeting cancer cells. The provided results are solely based on in-silico experimentation, and further experimental studies are required in future works to examine the effectiveness of the obtained results.

